# Activity-dependent remodeling of muscle architecture during distinct locomotor behaviors in *Caenorhabditis elegans*

**DOI:** 10.1101/2024.08.30.610496

**Authors:** Adina Fazyl, Akash Anbu, Sabrina Kollbaum, Andrés G. Vidal-Gadea

## Abstract

Muscle structure is dynamically shaped by mechanical use, yet how distinct locomotor behaviors influence sarcomere organization remains poorly understood. In *Caenorhabditis elegans*, crawling and swimming constitute discrete gaits that differ in curvature, frequency, and mechanical load, providing a tractable model for studying activity-dependent remodeling. Using confocal imaging of phalloidin-stained body-wall myocytes, we quantified myocyte geometry, sarcomere length, and sarcomere number across anterior, medial, and posterior regions in animals reared exclusively under crawling or swimming conditions. Quantification and hypothesis testing used linear mixed models that accounted for repeated myocyte measurements within animals, with interaction terms testing region-specific effects of locomotor condition after IQR-based outlier removal. Swimming produced characteristic remodeling of body-wall muscles. Myocytes elongated globally, while selectively thinning in the mid-body, reducing cell area by ∼13 % relative to crawlers. Shape metrics confirmed this shift: circularity declined at mid- and tail-regions and anisotropy increased by ∼2–3 units. Sarcomere architecture exhibited parallel remodeling. Average sarcomere length shortened across the body (−0.19 µm in head, −0.35 µm in mid-body, −0.20 µm in tail), while sarcomere number increased anteriorly and medially (+0.77 and +0.65 sarcomeres per myocyte). The medial region also showed a significant rise in sarcomere density, indicating tighter serial packing. These adaptations mirror functional compartmentalization predicted from gait kinematics and parallel fast-fiber remodeling observed in vertebrate muscles. The results indicate that *C. elegans* muscles adapt their contractile lattice to sustained mechanical demand, linking neural gait selection and mechanosensitive signaling to long-term structural plasticity. This work establishes *C. elegans* as a model for dissecting the conserved pathways that couple muscle use to cellular architecture and provides a foundation for future comparisons of healthy and diseased muscle remodeling.

**Short summary:** Muscle cells in *C.* elegans change their structure according to how the animals move. Worms that swim develop shorter, more densely packed sarcomeres and elongated body-wall muscles, while crawlers maintain longer, broader fibers. These adaptations enhance flexibility and power transmission for high-frequency motion, linking neural gait selection and mechanosensitive signaling to long-term remodeling of the contractile lattice.

## Introduction

Animals adapt their movement patterns to the physical properties of their environment, and muscles must structurally accommodate these different mechanical demands. In *Caenorhabditis elegans*, locomotion alternates between two distinct gaits: crawling on solid surfaces and swimming in liquid. Complementary behavioral and genetic analyses confirmed that transitions between solid and liquid evoke distinct locomotor programs rather than a single continuum of speeds (Pierce-Shimomura et al., 2008), establishing that crawl and swim represent bona fide gaits with different mechanical characteristics. Early work from our group demonstrated that these gaits are selected through aminergic modulation, with serotonin promoting the transition from crawling to swimming, and dopamine stabilizing crawling once animals return to land (Vidal-Gadea et al., 2011).

Crawling on agar imposes low-frequency, low-curvature bending on the body wall, whereas swimming in liquid requires higher-frequency undulations and greater medial curvature (Beron et al., 2015). Muscle-intrinsic mechanosensation has emerged as a key determinant of gait-appropriate force production (Kang et al., 2010; Wen et al., 2012; Thomas et al., 2024). Fazyl et al. (2024) showed that the mechanosensitive ion channel PEZO-1 localizes to the sarcolemma of body-wall myocytes, where it differentially modulates calcium dynamics and contractile output during swimming and crawling. Loss of PEZO-1 activity impairs swimming while modestly enhancing aspects of crawling, indicating that muscle mechanotransduction contributes to matching force generation to environmental load.

Beyond acute modulation of contractile activity, muscles remodel their internal architecture with chronic changes in use (Ferraro et al., 2014; Franchi et al., 2017; Pillon et al., 2020). In vertebrates, sustained mechanical loading alters sarcomere number, spacing, and lattice geometry to optimize contractile efficiency (Højfeldt et al., 2023). Whether comparable activity-dependent remodeling occurs in invertebrate muscles has remained largely unexplored. Our recent imaging studies revealed that the laterally interconnected sarcomere networks of *C. elegans* body-wall muscles remodel with mechanical demand and display structural failure in dystrophin-deficient mutants (Fazyl et al., 2025). These findings suggest that nematode muscles possess a dynamic, plastic contractile architecture capable of adapting to altered forces.

Here we focus on the healthy muscle response to the locomotor environment, isolating the effects of gait-related mechanical demand on myocyte morphology and sarcomeric organization. Using confocal imaging of phalloidin-stained myocytes from animals reared exclusively under crawling or swimming conditions, we quantified myocyte area, shape descriptors, sarcomere number, and sarcomere length across anterior, medial, and posterior regions of the body wall musculature. These measurements reveal coordinated, region-specific remodeling of the contractile lattice: swimming induces shorter sarcomeres, increased anisotropy, and higher sarcomere density, changes consistent with serial remodeling and redistribution of contractile material toward regions of greatest mechanical demand. Integrated with prior behavioral (Vidal-Gadea et al., 2011; Pierce-Shimomura et al., 2008) and mechanotransduction (Fazyl et al., 2024) studies, these findings link neural gait selection and PEZO-1-mediated feedback to long-term structural adaptation of muscle, positioning *C. elegans* as a comprehensive model for how environmental mechanics shape muscle function from neural control to cellular architecture.

## Materials and Methods

### Strains and maintenance

Wild-type *Caenorhabditis elegans* (N2, Bristol strain) were used in all experiments. Animals were maintained on nematode growth medium (NGM) agar plates seeded with *Escherichia coli* OP50 at 20 °C under standard conditions (Brenner, 1974). All assays used age-synchronized day-1 adults obtained by alkaline hypochlorite bleaching of gravid hermaphrodites followed by timed development on NGM plates.

### Locomotor regimens

To examine activity-dependent remodeling, animals were reared exclusively under one of two locomotor conditions from the L1 stage until the day of imaging. For crawling, worms developed on standard NGM agar plates seeded with OP50. For swimming, synchronized embryos were placed in 1.5 mL microcentrifuge tubes containing liquid NGM supplemented with OP50 (optical density approximately 0.8) and gently agitated on a nutator at 20 °C to maintain oxygenation (Laranjeiro et al., 2019). Animals were transferred to fresh liquid media every 48 hours.

### Sample preparation and fixation

Animals were washed three times in liquid NGM buffer, anesthetized with 10 mM sodium azide, and fixed in 4% paraformaldehyde in phosphate-buffered saline (PBS) for 15 minutes at 20 °C. Fixed worms were transferred to poly-L-lysine-coated slides and permeabilized using the freeze-crack method (Duerr, 2013). Following permeabilization, samples were blocked for 30 minutes in 1% bovine serum albumin (BSA) in PBS and stained overnight at 4 °C with iFluor 488-conjugated phalloidin (1:400; Cayman Chemical) to visualize filamentous actin. Slides were washed in PBS and mounted in ProLong Gold antifade reagent (Thermo Fisher Scientific).

### Confocal imaging

Fluorescent images were acquired on a Leica SP8 confocal microscope equipped with Lightning deconvolution (63×, NA 1.40 oil-immersion objective). For each animal, Z-stacks spanning entire body-wall myocytes were captured (0.17 µm step size, 15 to 30 slices). Maximum-intensity projections were generated in Leica Application Suite X (LAS X, v3.5.5.19976) and exported as 16-bit TIFFs. Imaging was performed from anterior, medial, and posterior regions corresponding to muscle cells 1–10, 11–18, and 19–25, respectively (Gieseler et al., 2017).

### Image analysis

Quantitative analysis was performed using Fiji/ImageJ (Schindelin et al., 2012). For each myocyte, the following parameters were measured: myocyte area, perimeter, Feret diameter, MinFeret, major and minor axes, aspect ratio, circularity, and solidity (using the Analyze → Set Measurements tool); average sarcomere length, measured as the mean distance between consecutive Z-line actin peaks within a single myocyte using the Plot Profile function; and sarcomere number, determined by counting Z-line intervals along the long axis of the same myocyte. Derived variables included sarcomere density (sarcomere number divided by cell area), serial sarcomere density (sarcomere number divided by Feret length), and anisotropy (Feret divided by MinFeret). All measurements were performed blind to locomotor condition.

## Statistical analysis

All statistical analyses were conducted in R (version 4.5.0 (2025-04-11)) using a custom analysis pipeline. Data processing and visualization were performed using the tidyverse and ggplot2 packages. Prior to modeling, data was pre-processed using a robust, group-wise outlier removal procedure. This was performed independently for each dependent variable. Outliers were defined within each experimental group (for each unique condition×region combination, such as “Swim-Head”). This group-wise approach prevents data from one group (a high-variance group) from biasing outlier detection in another (a low-variance group). For a given variable, observations falling outside 1.5 times the interquartile range (IQR) of their specific group were excluded from that variable’s analysis.

A robust, two-pronged hybrid modeling strategy was implemented to account for the hierarchical structure of the data (multiple measurements nested within each worm). This strategy ensured that the most statistically appropriate model was automatically selected for each dependent variable based on its specific data characteristics. The core model aimed to test the fixed effects of condition (Swim vs. Crawl), region (Head, Mid, Tail), and their two-way interaction (condition*region).

We first attempted to fit a Linear Mixed Model (LMM) using the lmerTest package, with worm specified as a random intercept: Dependent Variable∼condition*region+(1 | worm). This model was fit using Restricted Maximum Likelihood (REML). The LMM was considered the appropriate model only if it met several criteria: the model converged without errors; the model was not “singular,” defined as the variance of the worm random effect being estimated at or near zero (< 1e-8); and the pre-calculated Intraclass Correlation Coefficient (ICC) for the worm group was greater than 0.01, indicating non-negligible clustering. For variables with significant, well-estimated inter-individual clustering, this LMM was retained.

If any of the LMM criteria failed, the model was considered inappropriate. In these cases, the analysis automatically fell back to a standard Ordinary Least Squares (OLS) regression model: Dependent Variable∼condition*region. This fallback was triggered from variables with no detectable inter-worm variance, such as Maximum Feret and Sarcomere number (ICC ≈ 0).

P-values and confidence intervals were calculated using methods appropriate for the selected model. For LMMs, inference for the fixed effects was derived using the Kenward-Roger (KR) correction (ddf=“Kenward-Roger”). The KR method is considered the gold standard for small-sample LMMs, as it provides a denominator degrees of freedom (DF) approximation and, critically, adjusts the fixed-effect standard errors to reduce small-sample bias. This was successfully applied to all LMMs, as seen in the final results (DF_method=“Kenward-Roger”). For OLS Models, standard OLS p-values are invalid as they ignore the clustered data structure. Therefore, for all fallback OLS models, we calculated cluster-robust standard errors (using the sandwich and lmtest packages) by clustering on the worm identifier. This “sandwich” estimator provides valid p-values and confidence intervals that are robust to the non-independence of observations from the same worm, even if an LMM could not be fit. The specific correction used was “HC1” (DF_method=“ClusterRobustHC1”). “HC” stands for “Heteroskedasticity-Consistent,” and the “1” specifies a particular type of small-sample correction. This HC1 correction applies a degrees-of-freedom adjustment to the standard error calculation, which prevents underestimation of the variance and provides more reliable p-values when the number of clusters (in this case, worms) is small. All statistical tests were two-tailed, and a p-value < 0.05 was considered statistically significant.

Finally, to decompose significant interactions and directly test the effect of swimming versus crawling at each anatomical location, we conducted a post-hoc simple effects analysis using the emmeans package. We calculated the estimated marginal means (EMMs) for the “Swim vs. Crawl” contrast within each of the three regions (Head, Mid, Tail). This post-hoc analysis used a parallel modeling strategy. It fit the data to an LMM with Satterthwaite-approximated degrees of freedom, falling back to a standard OLS model if the LMM was invalid. This ensured the p-values for these specific contrasts were also robustly estimated and properly accounted for the data’s structure.

Raw data is provided as a supplement (Supplementary data 1). Full statistical analysis and the outlier log are provided in Supplementary data 2. For the R code used in the analysis please refer to Supplementary data 3.

### Data availability

All raw measurements and statistical outputs are provided as a supplement (Supplementary Materials) and figure source data. Representative imaging stacks and analysis scripts are available at https://figshare.com/s/8ea23d743d7c3e5740d0.

### Artificial Intelligence Tools

ChatGPT-4 was used to assist with manuscript editing and grammar checking. All scientific content, data analysis, and conclusions remain the original work of the authors.

## Results

### Experimental design and measurement strategy

To determine how sustained locomotor behavior influences muscle architecture, we compared body-wall myocytes from wild-type *C. elegans* reared exclusively under crawling or swimming conditions (Figure 1A). In *C. elegans* this is made possible by the geometry of the body wall musculature. Specifically, body wall myocytes occur in four quadrants and consist of single cells containing a singular layer of non-overlapping sarcomeres. This allows the use of maximal projections from confocal stacks to capture the entire contractile machinery within each cell without the confound of multiple sarcomeres overlapping in the z plane (Gieseler et al., 2017). Confocal imaging of phalloidin-stained animals revealed clear differences in myocyte geometry and sarcomere organization between swimming and crawling conditions (Figure 1B to D). Measurements were obtained from anterior, medial, and posterior myocytes (myocytes m1-10, m11-18, and m19-25, respectively). Each myocyte was outlined to determine geometric parameters (area, Feret, MinFeret, aspect ratio, circularity), and sarcomere number and length were quantified along the long axis (Figure 2A). Derived metrics included sarcomere density (number per area) and anisotropy (Feret divided by MinFeret). Our approach focused on myocyte area rather than volume. We decided to focus on this cellular dimension because it was the closest associated with the planar architecture of the contractile machinery.

**Figure 1.**
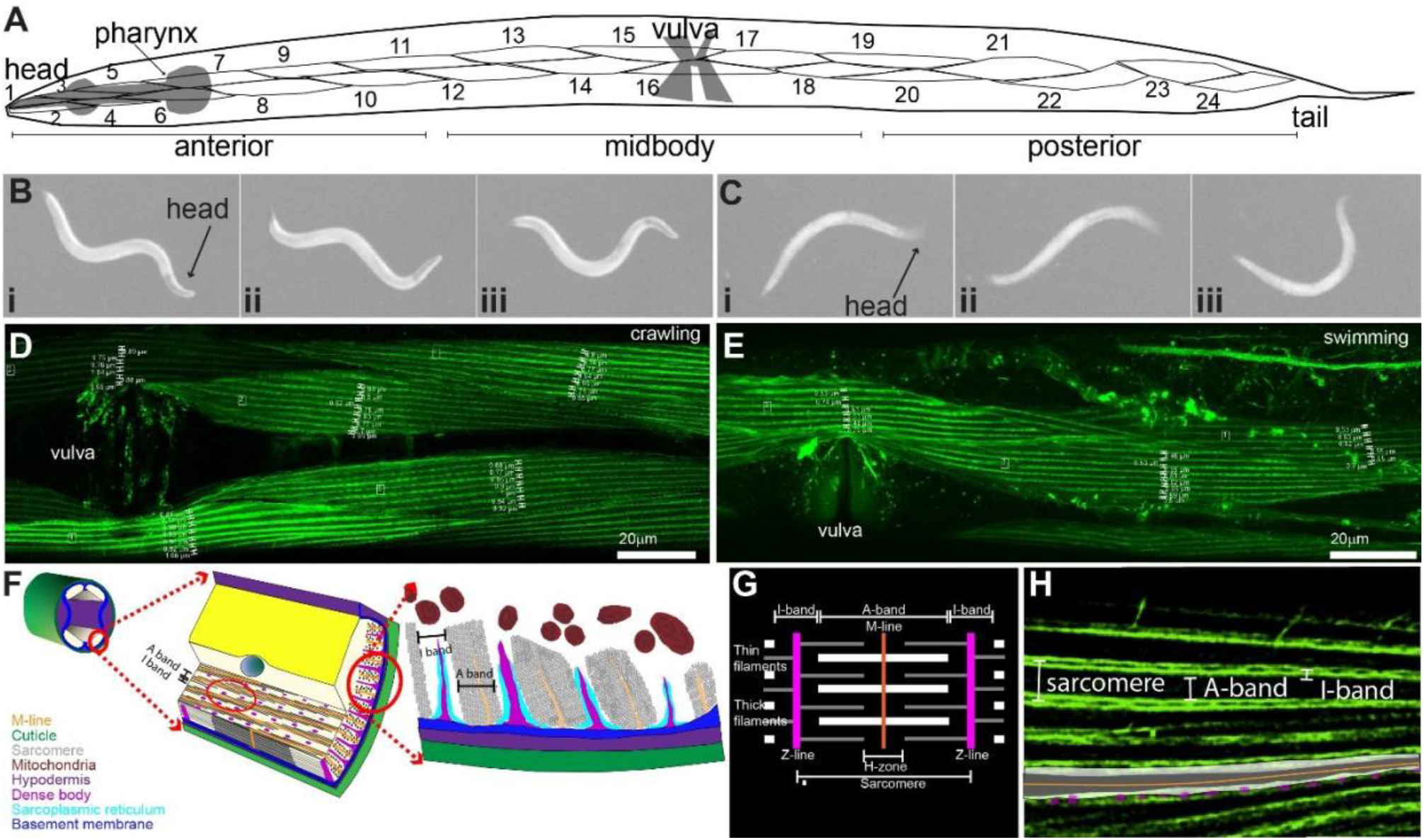
Experimental overview and measurement workflow. **A)** Schematic of *C. elegans* body-wall musculature showing anterior (head), medial (mid-body), and posterior (tail) regions analyzed. **B)** Representative bright-field stills of worms during crawling on agar (left) and **(C)** swimming in liquid (right), illustrating posture and curvature over time **(i–iii). (D–E)** Confocal micrographs of phalloidin-stained mid-body myocytes from crawling and swimming animals, respectively. Actin filaments delineate sarcomeres used for morphometry. Sample sarcomere measurements are displayed for four crawling **(D)** and three swimming **(E)** myocytes. **F)** Schematic diagram of *C. elegans* body-wall musculature styled after Gieseler et al. (2017). **G)** Typical sarcomere arrangement in mammalian striated muscle compared with analogous structures in *C. elegans* **(H)**.

**Figure 2.**
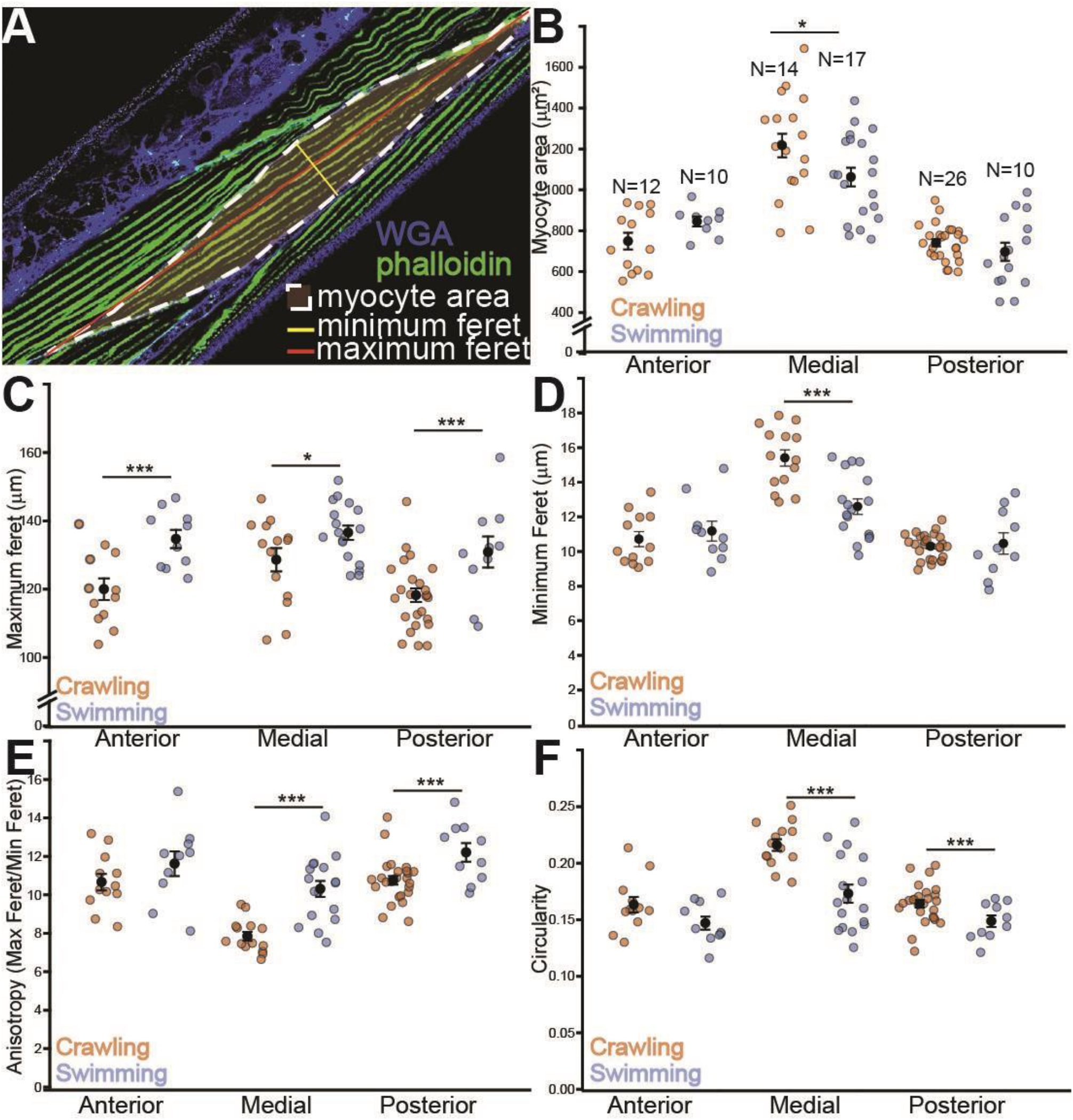
Myocyte geometry adapts to locomotor mode through global elongation and region-specific thinning. **A)** Representative confocal micrograph illustrating the quantitative analysis workflow. Body-wall muscle is stained with phalloidin (green) to visualize F-actin, and WGA (blue) stains cell boundaries. Overlays demonstrate the key geometric measurements taken for each myocyte: myocyte area (white dashed outline), maximum Feret diameter (red line), and minimum Feret diameter (yellow line). **B)** Area. Region-dependent effect of swimming (LMM, significant condition × region interaction, p=0.033), with smaller medial areas in swimmers (simple effect, p=0.032). (Ns: Head C=13, S=9; Mid C=18, S=20; Tail C=26, S=15). **C)**Feret (maximum diameter). Greater Feret in swimmers across regions (OLS-CR, main effect of condition, p=0.00019). (Ns: Head C=13, S=10; Mid C=18, S=18; Tail C=25, S=15). **D)** MinFeret (minimum diameter). Selective medial thinning in swimmers (LMM, significant condition × region interaction, p=0.0027). (Ns: Head C=13, S=8; Mid C=18, S=20; Tail C=26, S=16). **E)** Anisotropy (Feret/MinFeret). Increased anisotropy in swimmers, most pronounced medially (OLS-CR, significant condition × region interaction, p=0.015). (Ns: Head C=13, S=9; Mid C=18, S=20; Tail C=22, S=14). **F)** Circularity. Reduced circularity specifically in medial and tail myocytes of swimmers (OLS-CR, significant condition × region interaction; simple effects p<0.001 for Mid and Tail). (Ns: Head C=12, S=10; Mid C=16, S=20; Tail C=25, S=15). C=Crawl, S=Swim. Data are shown as mean ± s.e.m. N = 96–103 myocytes from 92–97 animals (total, depending on variable) after 1.5×IQR filtering.

### Myocyte geometry adapts to locomotor mode through region-specific thinning and global elongation

Swimming produced coordinated changes in myocyte geometry, driven by a combination of global adaptations and region-specific remodeling (Figure 2A-F). Across regions, myocytes elongated substantially: the mean Maximum Feret diameter increased from 118.2 ± 12.1 µm in crawling head muscles to 134.6 ± 8.5 µm in swimmers (p = 0.0006). Similar elongation occurred medially (from 127.1 µm to 136.3 µm, p = 0.014) and posteriorly (from 117.0 µm to 129.8 µm, p= 0.0006) (Figure 2C). In contrast, cell width narrowed selectively in the mid-body. Minimum Feret decreased from 16.0 µm in crawling worms to 12.8 µm in swimmers (p < 1×10−^6^), whereas anterior and posterior widths were unchanged (Figure 2D). Correspondingly, myocyte area in the medial region decreased by 154 µm^2^ on average (1217 ± 243 µm^2^ vs. 1063 ± 207 µm^2^; p = 0.032), whereas head and tail areas were unaffected (Figure 2B). These shape adjustments yielded a consistent morphological signature. Anisotropy rose from 8.0 to 10.8 in the medial region (p < 10−^6^) and from 11.5 to 13.8 posteriorly (p < 0.0001), reflecting stronger elongation (Figure 2E). Circularity showed a significant locomotion effect (*p* = 0.080 overall) modulated by region (*p* = 0.018), decreasing notably in both mid and tail regions (from 0.223 to 0.178, p < 10−^6^; from 0.168 to 0.139, p = 0.0003), confirming a shift toward more elongated and angular cell shapes (Figure 2F).

Together these data indicate that swimming promotes global myocyte elongation combined with selective mid-body thinning and shape reorganization, suggesting distinct mechanical loading patterns along the body axis.

### Sarcomere organization reflects coordinated activity-dependent remodeling

Swimming altered not only myocyte geometry but also the organization of the contractile lattice (Figure 3A-D). Sarcomeres were significantly shorter in swimmers: average lengths decreased from 1.62 µm to 1.43 µm in the head (p = 0.003), from 1.85 µm to 1.49 µm in the mid-body (p < 10−^9^), and from 1.57 µm to 1.36 µm in the tail (p = 7 × 10−^5^) (Figure 3A). This widespread shortening indicates a global remodeling program rather than localized contraction artifacts.

**Figure 3.**
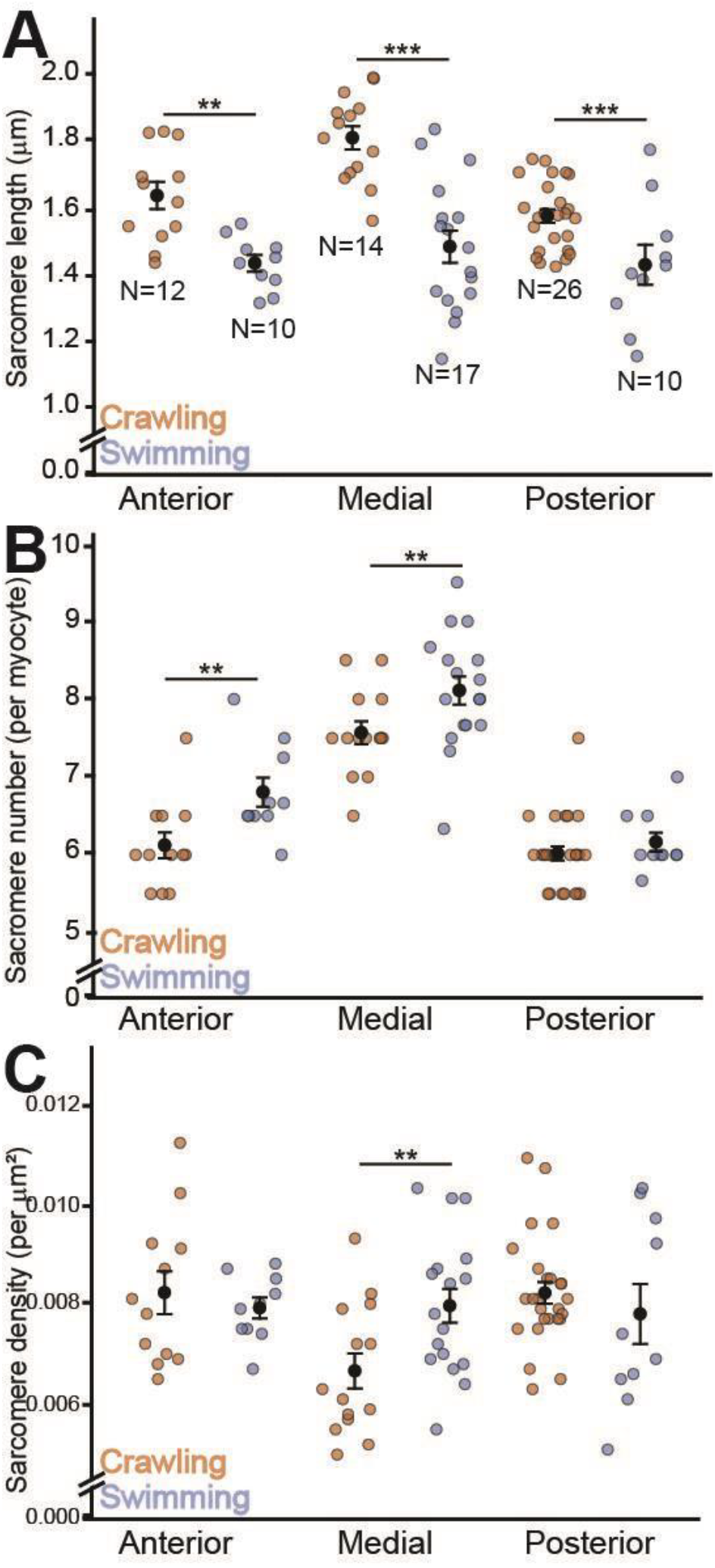
Sarcomere organization reflects coordinated activity-dependent remodeling. **A)** Sarcomere length. Sarcomeres are shorter in swimmers across all regions (LMM, main effect of condition, p=0.003). (Ns: Head C=13, S=10; Mid C=18, S=20; Tail C=26, S=16) **B)** Sarcomere number. Swimmers show higher sarcomere number in Head and Mid regions (OLS-CR, main effect of condition, p=0.0006), but this effect is absent in the tail (significant condition × region interaction for Tail, p=0.0019). (Ns: Head C=12, S=10; Mid C=15, S=20; Tail C=25, S=14) **C)** Sarcomere density (per µm^2^). Swimming significantly increases sarcomere density in the medial region (LMM, significant condition × region interaction, p=0.021). (Ns: Head C=13, S=10; Mid C=18, S=20; Tail C=22, S=16).C=Crawl, S=Swim. Data are shown as mean ± s.e.m. N = 96–103 myocytes from 92–97 animals (total, depending on variable) after 1.5×IQR filtering.

Swimming also increased sarcomere number (p = 0.0006), adding approximately 0.8 sarcomeres per myocyte in the head (p = 0.0013) and 0.6 in the mid-body (p = 0.0022), while the tail remained unchanged (p = 0.81) (Figure 3B). Sarcomere density showed a mild overall increase (p = 0.42) but a clear condition × region interaction (p = 0.021), driven by a medial rise from 0.0066 µm−^2^ to 0.0080 µm−^2^ (p = 0.010), suggesting serial addition and tighter packing (Figure 3C).

Integrating the geometric and sarcomeric data reveals a coherent myocellular adaptation program. During swimming, myocytes globally elongate, and the contractile lattice simultaneously shifts toward more, shorter sarcomeres. This coordinated remodeling is most pronounced in the medial body, where geometry (thinning and area reduction) and lattice organization (serial sarcomere addition) change together to optimize the myocyte for the high-frequency, high-flexibility contractions required for swimming, rather than the high-force output of crawling.

### Regional specialization and coordinated remodeling of cell size and sarcomere number

To test whether the geometric and sarcomeric adaptations documented above reflected coordinated changes within individual myocytes, we examined the relationship between myocyte area and sarcomere number. In anterior muscles, a modest positive correlation emerged when data were pooled across locomotor conditions (r^2^=0.33, p=0.005, n=22), suggesting that larger anterior myocytes accommodate proportionally more sarcomeres in series (Figure 4A). This relationship was not significant within crawling or swimming animals alone, indicating variability in how individual myocytes scale sarcomere number with size. In contrast, medial and posterior regions showed no significant correlation between area and sarcomere number (Medial: r^2^=0.05, p=ns, n=31; Posterior: r^2^=0.10, p=ns, n=36) (Figure 4B-C), indicating that sarcomere addition in these regions occurs independently of cell size changes.

**Figure 4.**
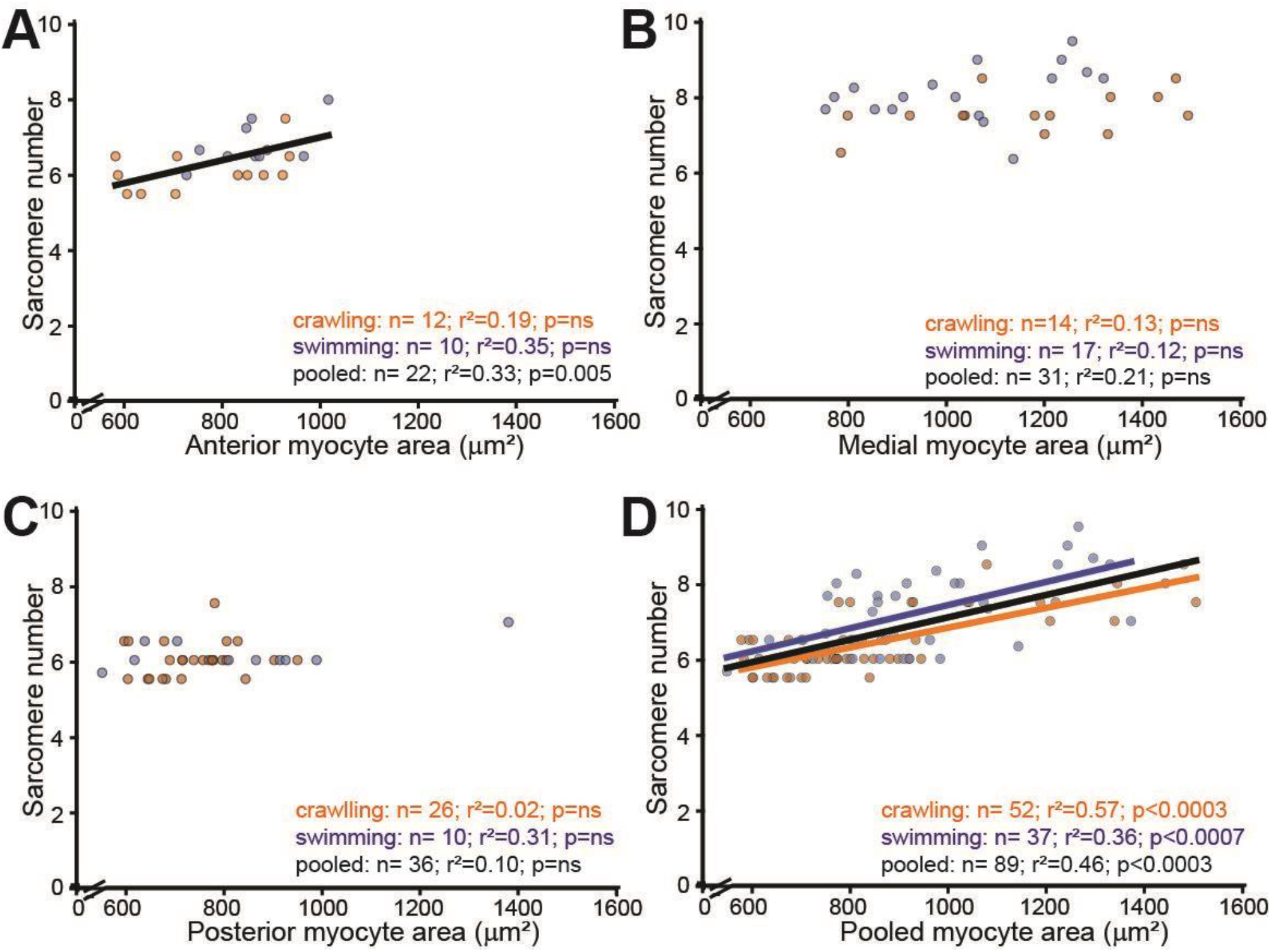
Myocyte area and sarcomere number show region-dependent scaling relationships. Scatter plots show the relationship between sarcomere number (x-axis) and myocyte area (y-axis) in (**A**) anterior, (**B**) medial, and (**C**) posterior body regions, with (**D**) all regions pooled. Within individual regions, correlations are either weak (anterior: r^2^=0.33, p=0.005) or absent (medial: r^2^=0.05, p=ns; posterior: r^2^=0.10, p=ns). When all regions are pooled, both crawling (r^2^=0.57, p<0.0003) and swimming (r^2^=0.36, p<0.0007) animals display significant positive correlations, reflecting primarily the intrinsic differences between body regions rather than tight within-region scaling. Data are individual myocytes after 1.5×IQR outlier removal (N = 89 total: Anterior n=22, Medial n=31, Posterior n=36). Lines represent least-squares linear regression fits (pooled in black; crawlers in brown; swimmers in blue). The parallel condition-specific slopes in panel D indicate that swimming shifts sarcomere number and area through the regional remodeling documented in Figures 2 and 3 without fundamentally altering the scaling relationship.

When all regions were pooled, both crawling (r^2^=0.57, p<0.0003, n=52) and swimming (r^2^=0.36, p<0.0007, n=37) animals displayed significant positive correlations between area and sarcomere number (Figure 4D). The strength of this pooled correlation reflects primarily the intrinsic differences between body regions rather than a tight within-region scaling relationship. Specifically, medial myocytes are naturally larger and contain more sarcomeres than anterior or posterior myocytes in both locomotor conditions, driving the overall positive association. The parallel slopes of the condition-specific regression lines indicate that swimming does not fundamentally alter how sarcomere number scales with area; instead, swimming shifts both parameters through the regional remodeling patterns documented in Figures 2 and 3.

Collectively, these data identify a distributed remodeling program. Swimming drives global myocyte elongation and sarcomere shortening across all body regions while simultaneously inducing dramatic region-specific changes in the mid-body, where muscles thin by approximately 3 µm in minimum diameter, lose approximately 150 µm^2^ of cross-sectional area, and increase sarcomere density. This compartmentalized response, combining uniform adjustments for contraction speed with targeted mid-body remodeling, aligns with the mechanical demands of swimming, where curvature and bending strain peak at the mid-body.

## Discussion

Our results show that locomotor context shapes myocyte architecture through coordinated and region-specific remodeling. Across the body, swimming drove elongation of myocytes and robust sarcomere shortening, coupled with increased sarcomere number in anterior and medial regions. These adjustments collectively tune the muscle lattice for high-frequency undulation and increased flexibility in liquid.

The most striking adaptations localized to the mid-body, where muscles thinned, reduced area, lost circularity, and increased anisotropy. This structural narrowing coincided with a rise in sarcomere density and a marked shortening of individual sarcomeres, forming a dense contractile lattice optimized for rapid curvature changes. Such compartmentalized remodeling aligns with mechanical analyses showing that curvature and bending strain peak medially during swimming, demanding both flexibility and continuous force transmission.

The correlation analysis between myocyte area and sarcomere number revealed that these two parameters scale together primarily at the level of regional identity rather than through tight within-myocyte coupling. Within individual body regions, the relationship was either weak (anterior) or absent (medial and posterior), yet when all regions were pooled, a strong positive correlation emerged. This pattern indicates that the remodeling program operates through region-specific set points rather than a simple rule linking cell size to sarcomere number. Medial myocytes, whether in swimmers or crawlers, are intrinsically larger and contain more sarcomeres than anterior or posterior myocytes, establishing distinct contractile configurations for each body segment. Swimming modulates these regional baselines, adding sarcomeres anteriorly and medially while selectively thinning the mid-body, without fundamentally altering the regional scaling relationships. This compartmentalized strategy allows independent tuning of different mechanical properties (force output, shortening velocity, strain resistance) along the body axis, matching contractile architecture to segment-specific demands during undulatory locomotion.

These findings fit with a mechanotransduction framework in which activity and load history tune muscle structure. Prior work from our group and others points to stretch-activated channels, including PEZO-1, as candidate sensors of mechanical state (Komandur et al., 2023; Fazyl et al., 2024). The medial-dominant remodeling observed here provides a testable anatomical substrate for such sensing. In future experiments, manipulating PEZO-1 in muscle and measuring the same geometric and lattice endpoints should reveal whether these channels are required to couple behavior to structural adaptation. More broadly, the pattern we document mirrors principles relevant to disease. In dystrophin deficiency, muscles face elevated mechanical stress and impaired force transmission (Petrof et al. 1993; Allen, Whitehead and Froehner, 2016). An adaptation program that increases sarcomere number while thinning and reshaping cells in regions of highest strain may fail or misfire without proper membrane stabilization, offering a route to early dysfunction.

Our approach has limitations that set the scope of inference and point to future work. First, we measured myocyte area rather than volume. We worked from maximum intensity projections of z-stacks spanning the entire myocyte layer. Because body wall muscle in this animal presents a single myofibrillar layer under the membrane, this approach likely captured the dominant activity-induced changes, yet it cannot exclude depth-wise remodeling. Future studies with volumetric reconstructions or light-sheet imaging could address this directly. Second, we focused on day one adults. This decision aligns with common practice and facilitates comparison with prior literature. However, these animals continue to grow and remodel over subsequent days. Extending the analysis through day five adults would determine whether the patterns we report are stable features of the adult program or evolve with continued growth. Third, we labeled F-actin with phalloidin to quantify sarcomeres. This marker is well suited for lattice geometry but does not report complementary components that may remodel with activity, including myosin heavy chain, α-actinin, costameric proteins, and membrane complexes. Parallel labeling or genetic reporters will be important to resolve which molecular compartments track with the geometric and lattice changes. Fourth, the environmental contexts are not identical beyond mechanics. Animals raised and assayed in liquid and on agar engage distinct behavioral programs and may experience different oxygenation, feeding, and sensory inputs (Vidal-Gadea et al., 2011 and 2012). We therefore cannot attribute every observed difference uniquely to mechanical challenge. What we can say is that worms develop and reproduce robustly in both environments, and the structural adjustments we measure are therefore physiologically relevant, even if multiple inputs contribute. Finally, one variable, circularity, required an ordinary least squares model with cluster-robust standard errors due to a near-singular random-effects structure after filtering. The direction and magnitude of the effect were consistent with the mixed-model results for the other variables, but this analytic caveat should be noted.

In sum, *C. elegans* body wall muscle exhibits a coordinated adaptation program that couples cell-level geometry to sarcomere-level organization, with the strongest remodeling in the medial body. This pattern provides a simple, quantitative framework for testing how specific mechanosensory pathways, including PEZO-1 (Komandur et al., 2023; Fazyl et al., 2024), and force-transmission scaffolds, including dystrophin-associated complexes, translate behavior into structural change.

## Supplementary Materials

**Supplementary File 1**. Spreadsheet with raw measurements used in this study

**Supplementary File 2**. Spreadsheet with descriptive and comparative statistics associated with the figures in this study.

**Supplementary File 3**. R code used for analysis.

## Author Contributions

A.F. conducted the experiments, performed imaging and quantitative analysis, wrote the manuscript. A.A. and S.K. assisted with data collection and analysis. A.G.V.G. conceived the study, supervised the research, and wrote the manuscript with input from all authors.

## Data Availability

All raw measurements, statistical outputs, representative imaging stacks, and analysis scripts are available at https://figshare.com/s/8ea23d743d7c3e5740d0.

## Acknowledgements

Some strains were provided by the CGC, which is funded by NIH Office of Research Infrastructure Programs (P40 OD010440).

